# TISMorph: A tool to quantify texture, irregularity and spreading of single cells

**DOI:** 10.1101/372755

**Authors:** Elaheh Alizadeh, Wenlong Xu, Jordan Castle, Jacqueline Foss, Ashok Prasad

## Abstract

A number of recent studies have shown that cell shape and cytoskeletal texture can be used as sensitive readouts of the physiological state of the cell. However, utilization of this information requires the development of quantitative measures that can describe relevant aspects of cell shape. In this paper we develop a toolbox, TISMorph, to calculate set of quantitative measures to address this need. Some of the measures introduced here are used previously and others are new and have desirable properties for shape and texture quantification of cells. These measures, broadly classifiable into the categories of textural, spreading and irregularity measures, are tested by using them to discriminate between osteosarcoma cell lines treated with different cytoskeletal drugs. We find that even though specific classification tasks often rely on a few measures, these are not the same between all classification tasks, thus requiring the use of the entire suite of measures for classification and discrimination. We provide detailed descriptions of the measures, well as TISMorph package to implement them. Image based quantitative analysis has the potential to become a new field of biological data (“image-omics”), providing quantitative insight into cellular processes.

## Introduction

The shape of a cell spread on a substrate is determined by the balance between the internal and external forces exerted on the cell boundary. The cell exerts forces and responds to external forces, from the extra-cellular matrix (ECM) or from neighboring cells, with the help of molecular motors and the cellular cytoskeleton, which is thus the ultimate determinant of cell shape [1, 2]. The cytoskeleton is a complex network, made of three major kinds of filaments – f-actin, microtubules and intermediate filaments – that form a cross-linked dynamic meshwork in the cytoplasm, providing shape and structure to the cell [1, 3]. The most dynamic constituent of the cytoskeleton, which is especially important in force generation and motility, is the filamentous actin (f-actin) network [4]. The f-actin network is directly involved in the formation of lamellipodia and filopodia through polymerization of f-actin against the cell membrane [5]. A third kind of cellular protrusions, blebs, are a result of the cortical actin network detaching from the cell membrane [6], and the convex shapes of adherent cells have been shown to result from myosin-II driven actin contractility [7].

The f-actin network is also deeply involved in force generation, force sensing and mechanotransduction. Contractile forces generated by myosin motors within cytoskeletal networks, membrane extension caused by actin polymerization, changes in osmotic pressure by opening of water or ion channels are examples of internal forces that play a role in shape of a cell. External forces leading to shape changes are applied through neighboring cells or ECM [8]. Actin filaments can generate and also resist mechanical stresses and cell deformation. But they can also eventually reorganize and change their structure, thereby sometimes relaxing external stresses. Different mechanical properties of the cell cytoskeleton and ECM will lead to different shapes for the cell. Thus the f-actin network is primarily responsible for the shape acquired by the adherent cell. It follows that the structure of the f-actin network must be related to the global shape of the cell, though the exact relation between the two is likely to be complex and non-linear.

Image-based screens have been used as a marker and predictor of cellular phenotype and behavior. Advancements in microscopy technology has provided the means to capture subcellular organization and cell shape with high resolution. However, our ability to gain insight into cellular processes through subcellular organization and cell shape is limited by the quantitative measures that we use to represent them. In machine learning algorithms information of each pixel in the image can be used to screen phenotype. However, implementing features of objects instead of pixels provides interpretable results at single cell resolution, which is more beneficial in biological applications. In addition using object features leads to reduced noise in the data, and may improve results.

Cell shape has been investigated as a marker and predictor of cellular phenotype and behavior. In our previous work, we used Zernike moments and geometric parameters as a measure of cell shape to distinguish between high metastatic and low metastatic osteosarcoma cancer cell lines with 99% accuracy [9, 10]. Other groups have also reported that cell shape can predict tumor grade [11], changes in the nuclear/cytoplasmic ratio of NFkB [12], and YAP (Yes-associated protein), [13], chemosensitivity of human colon cancer cell lines [14, 15], differences in motility [16], forms of motility [17], the progress of the Epithelial to Mesenchymal transition (EMT) [18] and the differentiation of Human Mesenchymal Stem Cells (hMSCs) into osteoblasts [19].

In addition to the importance of the actin cytoskeleton in determining the shape of the cell and the nucleus, the structure and organization of the f-actin network may provide additional information that can improve the prediction of cell physiology and better distinguish between cells in different states. There is a case to be made, therefore, of including measures of actin organization into studies of cell shape and its relation with cell phenotype, especially as actin staining is often used as the primary mean to determine the shape of the cell. Actin staining also provides textural information, which is directly related with actin structure, and as we show below, addition of this textural information significantly improves the discrimination between cell types.

The importance of cell and nuclear morphology and cytoskeletal organization as a tool for understanding and predicting cell behavior raises the need to appropriately quantify the shape and cytoskeletal texture of a cell. Here we introduce TISMorph as a tool to quantify Morphology and sub cellular structure of a cell. Good quantitative measures, should be capable of capturing biologically important differences between different experimental conditions. However, not all measures will be optimal for each comparison, and thus one will have to begin with a larger set of quantitative measures and discard uninformative ones if needed. Furthermore, good measures must not show significant differences between technical replicates in the same experiment. Based on these arguments we chose to test the measures calculated by TISMorph for cells treated with pharmacological modulators of the cytoskeleton. Features calculated by TISMorph can also be generalized to be used to quantify other subcellular structures such as intermediate filaments, plasma membrane, endoplasmic reticulum, mitochondria or even super cellular structures such as histopathology images and Magnetic resonance images of brain. These quantitative measures also meet the criteria proposed for good morphometrics by Pincus et. al. [20], i.e., that shape measures should possess fidelity, capture biologically important variation and be meaningful and interpretable.

## Experimental methods

For this study DUNN and DLM8 osteosarcoma cancer cells were used. The DLM8 line is derived from the DUNN cell line with selection for metastasis [21]. Therefore, DLM8 is closely related to DUNN except for degree of its invasiveness. Both cell lines were a gift from Dr. Douglas Thamm (Colorado State University, CO, USA). They were cultured on glass detergent washed and air dried (GDA) substrates with standard culture conditions of 37*°C* and 5% carbon dioxide concentration in Dulbecco’s Modified Eagle Medium (DMEM) (Sigma Alrdich). DMEM was supplemented with 10% EquaFETAL Fetal Bovine Serum (Atlas Biologicals) and 100 *Units/ml* penicillin with 100 *μg/ml* streptomycin (Fisher Scientific-Hyclone). After 45 hours of culture, cells were incubated with different cytoskeletal drugs with description, conditions, and vendors listed in the Table 1 for 3 hours. Then the cells were washed and fixed with 4% paraformaldehyde. Finally, they were fluorescently stained for nuclei (DAPI from Molecular probes) and actin (Acti-stain 488 phalloidin from Cytoskeleton, Inc). All the drugs were dissolved in Dimethyl sulfoxide (DMSO) (Fisher BioReagents^™^) and to drop its effect on cell shape and actin structure control study was also treated with DMSO with the same molarity as other drugs. Then the cells were imaged using fluorescent microscopy. Representative images of each cell line treated with these drugs are shown in Fig 1.

**Fig 1.**
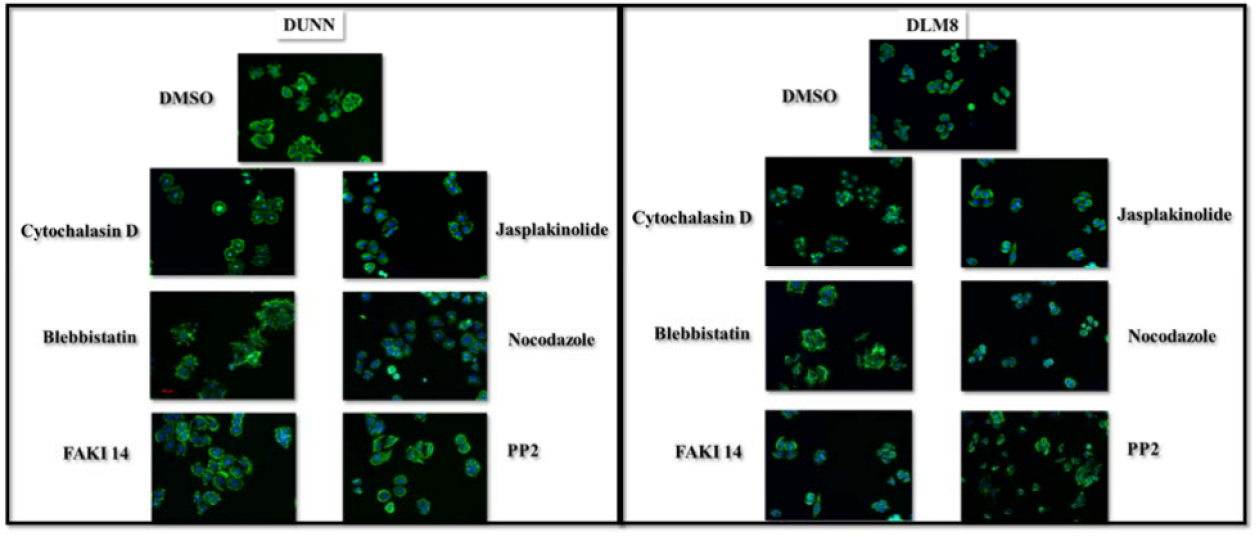
Representative images of DUNN and DLM8 osteosarcoma cancer cell lines with different drug treatments. Blue color represents nuclei and green color represents actin cytoskeleton.

## Image processing

In order to strike a balance between the throughput and the accuracy of the image processing, the image processing is fully supervised by the operator to reduce the number of artifacts and also automated as much as possible to speedup the processing of the images. The image processing code is stream-lined into a consecutive step-by-step workflow which consists of four steps as follows. 1) A graphic user interface (GUI) enables the binary thresholding of the actin and nucleus images. While a thresholding value is suggested automatically by Otsu’s method, users can easily adjust the thresholding value using a slide bar in the GUI by visually checking the original image and the thresholded image displayed side-by-side in the GUI. 2) Cell declustering in the thresholded images is done using an optimized template of the open-source software CellProfiler [22]. 3) The outputs from the CellProfiler are then visually examined by the operator and corrections can be made if necessary using the modules functionalized into this step. To facilitate the following analyses, each cell is centered and saved separately into one 1024×1024 image. Except for CellProfiler, all the other image processing codes were programmed in-house using Matlab (Mathworks) and are available on https://github.com/Wenlong-Xu/Image_Processing_Cell_Shape. Also available are the detailed protocol on how to configure the image processing codes and the CellProfiler template used for cell declustering.

## Data analysis methods

### Quantifying changes in multidimensional shape space

Each cell can be represented as a point in multidimensional shape space. For ease of analysis this vector can be projected down to a lower-dimensional space of the first few principal components (PCs). Differences between different conditions or different cell lines can be represented by p-value between their shape distributions for each PC space. While doing multiple comparisons, we will often pick the most informative principal component for our analysis from among the first four PCs. This is done in the following manner. We pick the PC whose worst case, i.e. largest p-value among all comparisons, is better (i.e. smaller p-value) than that of any other PC, so that it is the best single measure for distinguishing between all the comparisons. In other words, for each principal component maximum p-value between all comparisons is calculated as Eq (1).

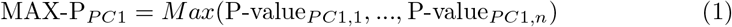

Where P-value_*PC1,n*_ is the p-value for *n^th^* comparison in the first principal component. Then from the first four PCs, the one that has the smallest MAX-P, maximum p-value between different comparisons, is chosen. In each shape category, we choose the best principal component by this criteria, and will describe them as the Primary Principal Components (PPC) of each shape quantification measure or category, for each analysis.

### Pearson correlation

Pearson correlation coefficient is calculated between features as shown in Eq (2). Here *r* is Pearson correlation coefficient, *x_i_* is the feature *x* for *i^th^* sample, *y_i_* is the feature *y* for *i^th^* sample, *μ_x_* is the average value of the feature *x* for all of the samples, and *μ_y_* is the average value of the feature *y* for all of the samples. This coefficient is calculated for all 14 cell lines and drug combinations (2 cell lines x 7 drugs) and the averaged coefficient is recorded. The results are shown in heat map plots in S1 Fig. In these plots the diagonal elements are correlation between features with themselves, so they are all 1. Also, Pearson correlation coefficient is symmetric so each heat map plot will be symmetric as well.

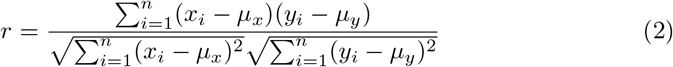

## Results

### Developing shape quantifiers

The actin structure is obtained from pixel intensities of labelled actin, which we can represent by a grayscale image, Fig 2a. These grayscale images of cells, or images of stained nuclei, are converted into a binary image (Fig 2c) and the coordinates of the edges are recorded (Fig 2b), yielding the cell boundary. To extract features of the morphology from this numerical data, we introduce three classes of shape measures, described as textural, spreading, and irregularity classes. Textural measures are divided into three subcategories called band based measurements, gray scale fractal dimension, and gray scale measures which are calculated based on the intensity plots of actin, Fig 2a. Spreading measures, which include measures based on a Zernike moment representation of shape as well as subcategories involving basic geometric parameters such as area and perimeter, are extracted from the binary representation of the cell (Fig 2c), its convex hull, or a similar image of the nuclei. Irregularity measures include parameters such as the waviness and roughness, which use information regarding the pixels at the boundary of the cell, Fig 2b. Each of these shape categories have been described below.

**Fig 2.**
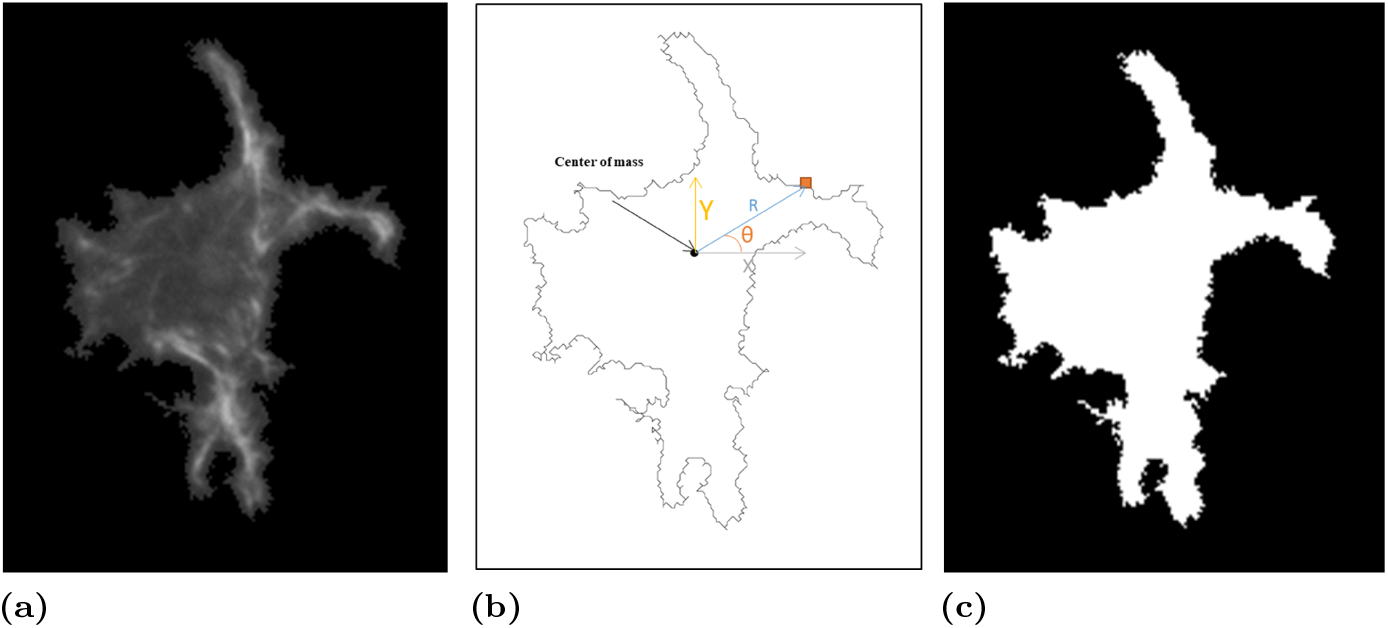
Representation of cell shape. a). Gray scale images using labeled actin. Intensity of each pixel is recorded and represented as a grayscale intensity plot. b). 2D outline: Position of each pixel in the boundary is recorded in polar or Cartesian coordinate. c). Binary image of actin.

#### Textural measures

##### Band based measurements

This parameter is sensitive to changes in the radial symmetry of the distribution of actin filaments. We divide the image into 10 equally spaced concentric regions (Δ*r*) around the center of mass of the image, called bands, as shown in S2 Fig. Five quantifiers are used to measure the differences in actin distribution between these bands. First, average intensity for each band is calculated. Then indices of the bands which have the lowest or the highest average intensity and their value are recorded. The last measurement is what is called above average adjusted intensity of the bands which is formulated in S1 Table along with other band based measures [23]. Band based measures are especially important in characterizing the actin distribution of the cells treated with Cytochalasin D, where actin has very unique symmetrical distribution. In these cells, dense foci are formed around the nucleus. The bands located in the central region of the cells are void of actin. At the outer bands, short linear actin structures aligned along the radius are observed.

##### Gray scale fractal dimension

Fractal dimension (*FD*) is a measure of roughness of objects, and can be applied to the characterization of texture in engineered and natural images. There are many methods to calculate this measure, and we chose the box counting method for its inherent simplicity. In this method the binary image is first covered with an evenly spaced grid with side length of *∈*. Then, the number of boxes which cover the fractal image are counted. This process is repeated by decreasing the side length and the *FD* is calculated based on Eq (3)

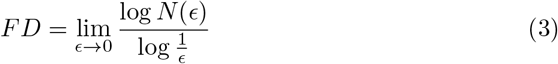

Here *∈* is the size of each box in the grid and the variable *N*(*∈*) is the number of the boxes which contain the fractal. We calculate *FD* based on a binarization of the grayscale image using edge detection methods to identify actin voids. This is done using four different edge detection methods in Matlab. To provide an example, binarized images of a cell from DUNN cell line treated with Cytochalasin D using these four edge detection methods are shown in S3 Fig. We found that different edge detection methods pick up different aspects of the actin structure, and so for every cell their gray scale image is binarized using these four methods, and their *FD* calculated [24].

##### Other gray scale measures

Haralik et, al introduced a procedure for quantifying the texture of satellite images based on the spatial relation between the gray tone of neighboring pixels in an image, for image classification [25]. This method is based on the Gray Level Co-occurrence Matrix (GLCM, sometimes called Spatial-Dependence Matrix), calculated as follows. For an image that has intensities of 1, 2,…, *g*, the co-occurrence matrix is a *g* x *g* matrix such that its ijth element is the number of the times that a pixel in the image has intensity equal to “i” and the pixel at a pre-defined distance 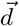 (which we choose to be 1 pixel) from it has intensity of *j*. An example of a 4 pixels x 4 pixels image with 5 gray tone levels is shown in S4 Fig. In a GLCM matrix when there are very few dominant transitions in gray tone of an image in neighboring pixels, the matrix will have small value for all of the entries. Each diagonal element of a GLCM matrix represents the number of the times that gray tone does not change in the neighboring pixel. Large diagonal elements imply that the image is homogeneous. Large numbers in the far upper right and far lower left of the matrix implies large transitions in intensity and high contrast in an image. After calculating the GCLM matrix, shown for the example image in S4(c) Fig., 23 different measures are calculated to quantify the texture in an image. The list of measures are tabulated in S2 Table [25-27]. For example one of the parameters is the contrast. For interpretation of this measure, it is useful to keep in mind that the contrast in an image is proportional to the changes in gray tone, *n* = |*i* − *j*|, so the far upper right and far lower left which have bigger value of *n* will have bigger contribution to the contrast parameter. For homogeneous images, the diagonal entries will be large, and will have a bigger contribution to the homogeneity measure. In this study we calculate the GLCM matrix for the vector 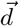 equal to 1 pixel in magnitude and with directions of 0*°*, 45*°*, 90*°*, and 135*°* and then the average value for each parameter is reported. This process yields rotation invariant measures.

#### Irregularity of boundary measures

In addition to cell spreading, irregularities in the boundary also carries information about the state of the cell. For example an irregular border could arise due to a large number of filopodia in the cell, which is a signature of a highly dynamic cytoskeleton. A highly contractile cell may retract from focal adhesions at the boundary creating many membrane protrusions that increase variability. To quantify irregularity of the cell boundary, the 2D boundary of the cell is used as shown in Fig 2c. We use the pixel positions that mark the boundary of the cell to calculate waviness, which estimates the periodic variation in the boundary, and roughness, which measures the non-periodic variation in the boundary, as discussed below.

##### Waviness measures

Using a Fourier series, a signal can be expanded in terms of linear combinations of orthogonal basis functions of sines and cosines with increasing frequencies. In the Fourier series expansion, it is assumed that the input signal is periodic. The outline of a cell is a closed curve, hence it is counted as periodic signal. The boundary of the cell could be represented in Cartesian coordinates, *x* and *y*, or polar coordinates, *ρ* and *θ*. Regardless of coordinate selection, it will lead to two independent signals which can be separately written as linear combination of cos and sin basis functions as Eq (4).

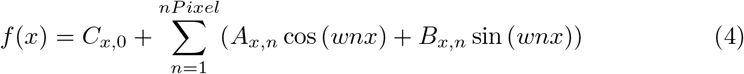

Where *nPixel* is the number of the pixels in the boundary of a cell and 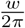 is fundamental frequency which is equal to 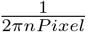. The variable *n* is an integer number and *n* * *w* is the *n^th^* frequency in the decomposition. Variables *A_x,n_* and *B_x,n_* are amplitudes of *n^th^* frequency, *C*_*x*,0_ is the mean of the signal, and *f*(*x*) is the input signal. If we use a very large number of frequencies, we can reconstruct the cell almost perfectly, but the number of descriptors will be unnecessarily high. There is a tradeoff between the number of frequencies (shape descriptors) used and the accuracy of the reconstruction. We are interested in quantifying shape without dealing with high number of parameters. Since the amplitude decreases with increasing frequency, we can filter higher frequencies and just use lower frequencies to decrease the number of descriptors used for the analysis. Here, with qualitative analysis of reconstructed cells with different frequency we decided to use only the first 35 frequencies as descriptor of the cell. Reconstruction of the shape of even rather irregular cells using these 35 frequencies is excellent, as shown in Fig 3. Finally, a remaining issue for the Fourier coefficients is that they are not rotationally invariant. We can construct a rotationally invariant measure by using Eq (5), which removes the phase difference in the Fourier expansion. This also reduces the number of parameters by half. Even though reconstruction of cell shape will not be possible with the rotationally invariant measures, our results show that they are nevertheless useful parameters for distinguishing between cells with different degree of variations in the radius. We will refer to these parameters as Waviness parameters. Interestingly, for Fourier parameters, all the features show low correlation coefficient with each other (0.35), suggesting that each coefficient carries mostly independent information. This observation was expected since we use orthogonal basis functions.

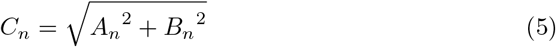

**Fig 3.**
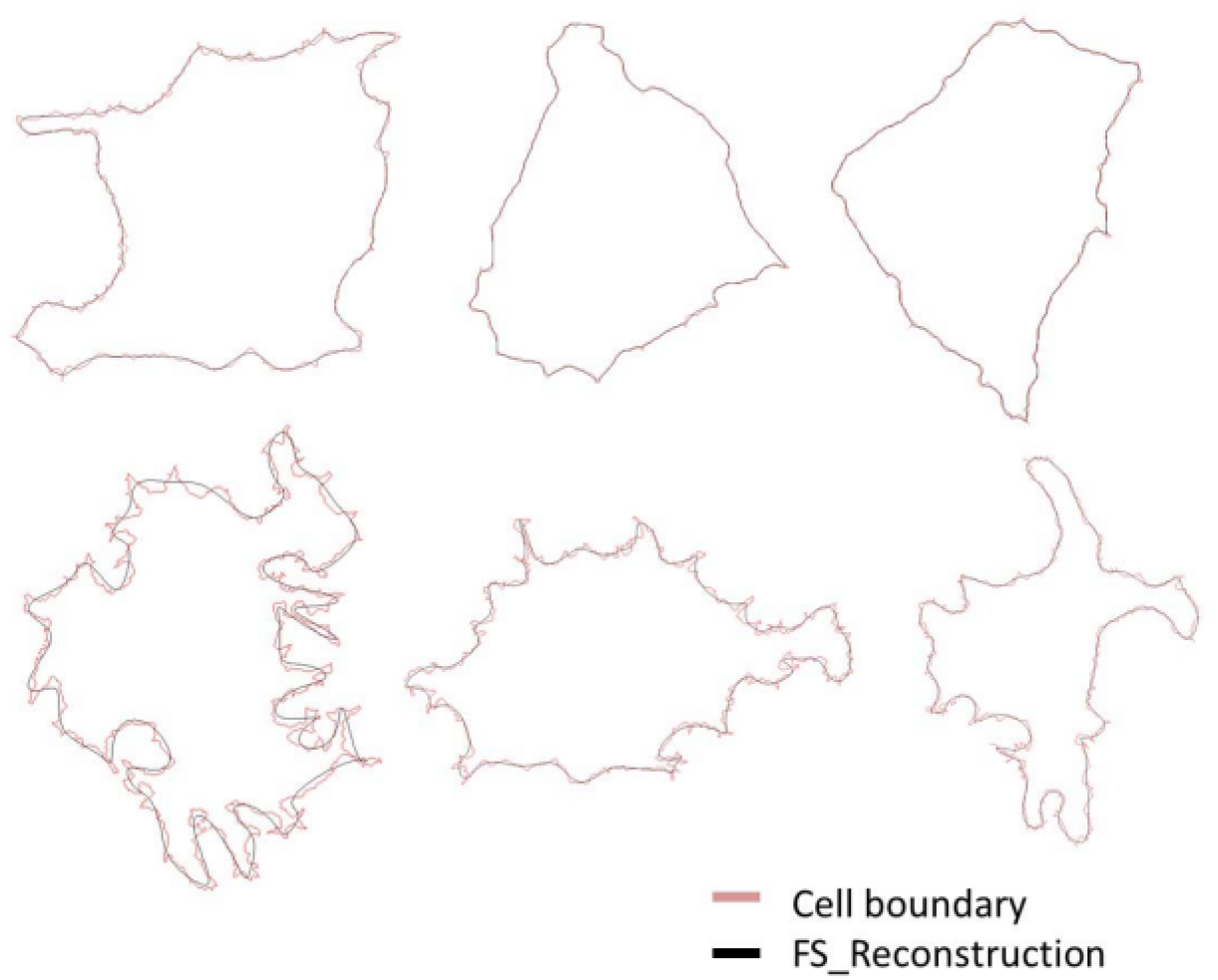
Reconstruction of the cells using Fourier decomposition. The red line is the actual boundary of the cell and black line is the reconstruction of the cell using the first 35 terms in the Fourier series expansion.

##### Roughness

As explained earlier, in the Fourier series decomposition of shape, higher frequencies have small amplitude and we neglect them. However, we can still obtain information about small amplitude variations from those high frequency terms. Following the method of Villanueva et. al. [28] to account for high frequency measures, we reconstruct the cell with the 35 Fourier components and subtract it from the original signal. The difference represents the roughness of the perimeter on a smaller scale than the variations picked up faithfully by the 35 Fourier frequencies. Statistical measures of the magnitide of this difference are called roughness measures and are listed in S7 Table. Visual examples of roughness as defined here can be seen in the lower panel of Fig 3.

#### Spreading measures

In this class of shape quantifiers, the binary image of cell and nuclei is used. These measures include geometric parameters for nuclei and cell, convex hull measures and Zernike moments for cells.

##### Geometric measures

Parameters like area, perimeter, major axis of fitted ellipse, minor axis of fitted ellipse, and their ratio are examples of geometric measures. All geometric measures used in this study to quantify cell and nuclei’s shapes are listed in S4 Table and S5 Table.

##### Convex Hull measurements

A convex polygon is a closed polygon such that the connecting line between any pair of points inside the polygon, lies completely inside the polygon. S5(a) Fig and S5(b) Fig. shows an example of a convex and non-convex polygon. The convex hull of a 2D shape is the smallest convex polygon that encloses the whole shape [29]. An example of the convex hull of a 2D cell shape is shown in S5(c) Fig. Convex hull geometric measures used in this study are listed in S6 Table.

##### Zernike moments

To calculate Zernike moments, the image of the cell is projected to the Zernike Polynomial basis function and the coefficients, called Zernike moments, are used as descriptor of cell shape. Zernike moments are complex numbers whose magnitude is rotation invariant, and can be made displacement invariant by moving the cell to the center of the imaging frame such that the center of mass of the image falls on the geometric center of the frame. The procedure has been discussed extensively previously [9],and Zernike moments have been used previously in other types of biological image recognition [30]. The Zernike moment expansion is also a basis function expansion like the Fourier series, and is capable of perfect reconstruction of the cell in principle. However in practice numerical errors prevent an accurate reconstruction when too many moments are used, and thus Zernike moments cannot capture the fine features of cell shape as well as Fourier series can. We found the best results when using about 147 Zernike moments (order upto 30, repetition for each order upto 10), which is what is currently calculated by TISMorph. These numbers can be changed by the user if needed.

### Validating performance of developed measures on osteosarcoma cells perturbed with different cytoskeletal drugs

#### Drugs which directly perturb microtubules, myosin-II or actin leave a unique signature on actin structure which is quantifiable by textural measures

We found that the most dramatic and unique effects on cell shape and texture arise from drugs that directly perturb microtubules or actin filaments. As shown in Fig 1, control cells acquire a polygonal shape on surfaces. Depolymerizing actin using Cytochalasin D leads to rounded cells where the actin has re-organized into alternating concentric rings of high and low density. Within each ring the actin is structured in very unique radial streaks! Increasing the depolymerization rate of microtubules using Nocodazole leads to a small rounded morphology. Inhibiting myosin II activity and decreasing cell’s contractility using Blebbistatin leads to increase in irregularities in the cell boundary. These changes in shape and structure of a cell are similar for both DUNN and DLM8 cell lines. Changes in structure and morphology of the cells using the other drugs in our experiments, with more indirect effects on the cytoskeleton, are subtle and not easily identifiable by eye. To quantify changes in shape and structure of the cells perturbed with different drugs, gray scale and binary image of the cells along with information of the boundary of the cells were used to measure all features in the 9 shape categories detailed above for each cell. S9 Table demonstrates quantified changes in all the measures for all the drugs. As demonstrated in this table the changes in textural measures for all the drugs are significant. This means that all the drugs that perturbed actin either directly, or indirectly through perturbing other cytoskeletal components, lead to significant changes in actin cytoskeleton organization.

#### Actin reorganization changes the spreading measures for both the cell and the nucleus

To explore changes in 2D shape of a cell and nuclei by changing actin distribution we compare the PPC of the geometric measures of the cell and nuclei. As shown in Table 2 perturbing actin significantly changes cell geometric measures other than for DLM8 cell line treated with Jasplakinolide. Interestingly changes in actin structure not only changes cell geometric measures, but it also changes nuclei geometric measures for all conditions other than both cell lines treated with Blebbistatin and DUNN cell line treated with Jasplakinolide and PP2 drugs. Zernike moments and Convex Hull parameters are similar in some respects. Although their quantification method is very different, as shown in reconstruction of the image of cells using Zernike moments (S6 Fig) both measures ignore the irregularities and fine fluctuations in the boundary. As demonstrated in Table 2, changes in the PPC for both measures are not distinguishable for DUNN cell lines treated with PP2, and both cell lines treated with FAKI 14.Moreover, hull geometric measures do not change significantly for DLM8 cells treated with Cytochalasin-D and DUNN cells treated with Jasplakinolide. In addition, Zernike moments do not change significantly for the DLM8 cells treated with Blebbistatin and DUNN cells treated with Cytochalasin-D. All other drugs lead to distinguishable changes in convex hull and Zernike moment measures. Since many cell shape parameters are expected to be correlated with each other, we performed a correlation analysis of all the measures within each shape quantification category, which are shown in S1 Fig. These results indicate that Zernike moments, cell geometric measures, nuclei geometric measures, and convex hull geometric measures are highly correlated with each other.

#### Cytoskeletal reorganization leads to changes in irregularities of a cell’s boundary

As shown in Table 2, waviness measures for all the drug conditions, except for DUNN cell lines treated with Jasplakinolide or PP2 and both cell lines treated with FAKI 14, changes significantly with respect to controls. In addition, the roughness measures change significantly for both cell lines treated with Blebbistatin, Cytochalasin-D, FAKI 14, and Nocodazole. In both cell lines, Jasplakinolide does not change the roughness measure significantly, which is also the case for the DUNN cell lines treated with PP2.

#### Different categories of shape quantifiers represent partly non-redundant shape information

While we have shown that each of our major categories and subcategories serve as good shape measures, in that they can be used to look for interpretable shape changes in different drug conditions, it is not clear whether we need all of them to represent shape. In order to estimate the degree of redundant information carried by the different shape categories, we calculated the Pearson correlation coefficient between all the features from different shape categories (shown in S1 File). In general, features from two different shape categories are relatively weakly correlated (< 0.4) except for Convex Hull and Cell Geometric features which are highly correlated with each other. There are a few other specific exceptions for which the features are also highly correlated. They are as follows. Zernike moments with *n* < 16 and *m* = 0 are highly correlated with Area. The correlation coefficient between Cell Area and Zernike Moment 0-0 is 1. It decreases with increasing order till it is almost zero for Zernike Moment (22-0), then it becomes negative and increases in magnitude (S7 Fig). The coefficient *C*_0_ from waviness features has a correlation of 1 with the Mean Cell Radius. It is also highly correlated with Cell Area (0.93) and Convex Hull area. Since the Fourier decomposition is based on a radial representation of the cell shape, *C*_0_ is a measure of the average cell radius, and should be expected to show these high correlations. Fractal Dimension measures also have high correlation with Cell Geometric, Nuclei Geometric, Gray Scale measures, and Zernike moments with *m* = 0 and *n* < 17. However, apart from these specific cases, the relatively high number of weak correlations between different shape categories implies that these shape categories contain non-redundant information about cell shape. Thus, quantitative shape analysis should ideally be carried out with representations from all of these shape categories in order for most efficient discrimination between different experimental conditions.

## Discussion

In this paper we introduce and provide the TISMorph package to quantify cell shape and cytoskeletal structure based on two dimensional images of cell morphology and actin structure. The Matlab toolbox used to process the images and quantify the shape and structure of a cell is shared in GitHub repository to be used by others. These toolboxes can be found in the following addresses, https://github.com/Wenlong-Xu/Image_Processing_Cell_Shape, https://github.com/Elaheh-Alizadeh/Quantifiction-of-shape-and-structure. Some of the textural and morphological features calculated by TISMorph are similar to the measures calculated by CellProfiler(CP) [22]. Geometric features of nuclei and cell, and gray scale measures calculated in this paper overlap with the measures calculated in CellProfiler by MeasureObjectSizeShape and MeasureTexture module. CP also calculates Zernike moments, though only to order 10,while we use Zernike moments up to order 30, because we find that fewer orders do not resolve objects sufficiently. Some of the measures we calculate are not implemented in the version of CP current during the time of submission. These are Fractal dimension, hull geometric measures, band based measures, and irregularity measures. However CP does calculate a few additional measures in the MeasureObjectIntensity module, which are statistical measures of intensity of objects such as mean and standard deviation of intensity, that are currently not included in TISMorph. Thus TISMorph includes significant additions to the previous state of the art as represented by the quantitative metrics calculated by CP.

The quantitative measures in the TISMorph package have been developed on the basis of empirical investigation of statistically significant differences between experimental conditions in principal component space. In this paper we explored the capacity of these quantitative measures to capture biologically important information. We perturbed the cytoskeleton of the cells with different drugs and explored their effect on cell shape and its structure. We first used textural measures to verify that the actin structure of the cell changes in the cells treated by cytoskeletal drugs and then we explored changes in the cell’s and nuclei’s 2D shape and irregularities of the cell boundary accompanied by changes in actin structure. Here we showed that most of the drugs used in this study directly or indirectly lead to significant changes in actin structure. Then we explored effect of changes in the structure of the cell on the measures of irregularities of cell boundary, nuclei spreading, and cell spreading. In most of the cases changes in actin structure are accompanied with significant changes in irregularities of cell boundary, and cell and nuclei spreading. The results show that textural measures and spreading measures are related but their relation is not simple, and the two classes of measures carry non-redundant information. It is worth mentioning that although we implemented textural measures to quantify actin structure, they can be used to quantify other sub-cellular and super-cellular structures as well. Shape quantification methods presented in this paper will prove useful for morphological screening for use in computer aided diagnostics in diseases such as cancer that are associated with cytoskeletal perturbations, assessment of qualitative cellular changes in different experimental conditions, and for mechanistic understanding of the determination of cell shape. In particular, morphological screening is emerging as a new high-throughput technique with wide applications in assessing functional biological responses [?, ?], and TISMorph should help increase the sensitivity and specificity of morphological comparisons.

## Supporting information

