## Supplementary material for "TISMorph: A tool to quantify texture, irregularity and spreading of single cells"

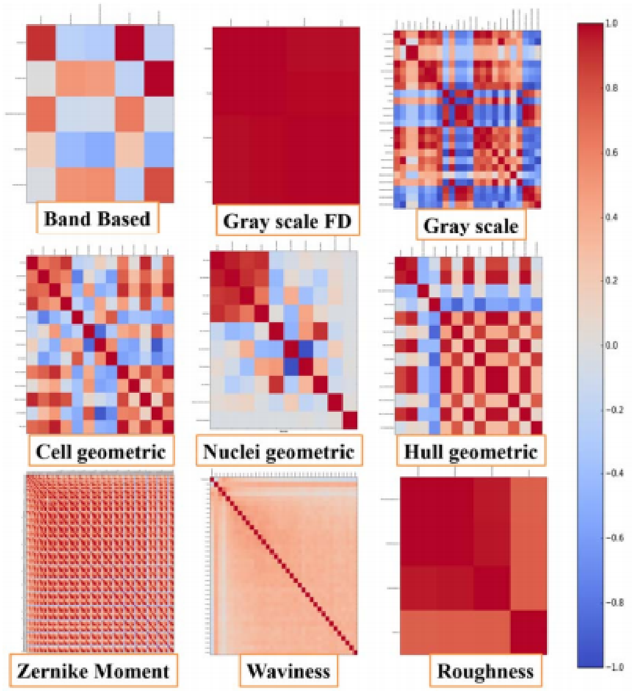


Supplementary Figure S. 1: Pearson correlation coefficient within each shape category. Within each shape category features are highly correlated other than band based, waviness (Fourier), and some of the nuclei measures. Each pixel is the average of Pearson correlation for 14 cases, 7drugs x 2 cell lines.

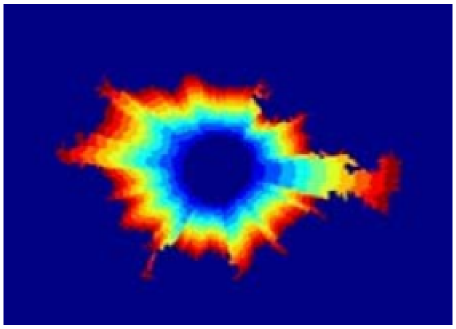

Supplementary Figure S. 2: Band based division of textural image of a cell. Cell is divided into 10 equally spaced radial bands. All the measures calculated based on this representation is listed in Table S1.

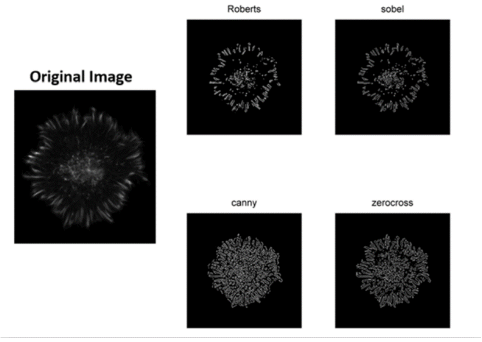


Supplementary Figure S. 3: Matlab Edging methods used to binerize gray scale image of actin. Four different methods of Roberts, Sobel, Canny, and zerocross are used in this paper.

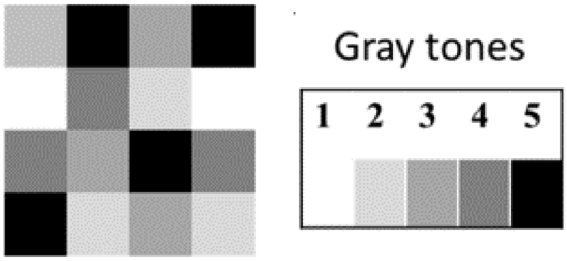


(a) (b)

| | | | |
|---|---|---|---|
| 2 | 5 | 3 | 5 |
| 1 | 4 | 2 | 1 |
| 4 | 3 | 5 | 4 |
| 5 | 2 | 3 | 2 |

(c)

| Gray tone of : origin↓ ,destination→ | 1 | 2 | 3 | 4 | 5 |
|---|---|---|---|---|---|
| 1 | (1,1) | (1,2) | (1,3) | (1,4) | (1,5) |
| 2 | (2,1) | (2,2) | (2,3) | (2,4) | (2,5) |
| 3 | (3,1) | (3,2) | (3,3) | (3,4) | (3,5) |
| 4 | (4,1) | (4,2) | (4,3) | (4,4) | (4,5) |
| 5 | (5,1) | (5,2) | (5,3) | (5,4) | (5,5) |

(d)

Supplementary Figure S. 4: Calculation of GCLM matrix. a) A cartoon of a 4pixels x 4pixels image. b) As shown in this figure the image has 5 gray tones. c) Numerical representation of image based on gray tone is shown in this figure. d) GCLM Matrix is 5 x 5 element and its $ij^{th}$ element is number of the times which pixel with intensity of $j$ is in distance $\vec{d}$ from the pixels with intensity of $i$. The vector $\vec{d}$ is an arbitrary vector but should be defined in advance. In this paper $\vec{d}$ is defined to have length of one and it is defined in 4 directions of $0^o$, $45^o$, $90^o$, and $135^o$. Gray scale measures are calculated based on four directions and their average is reported in this paper.

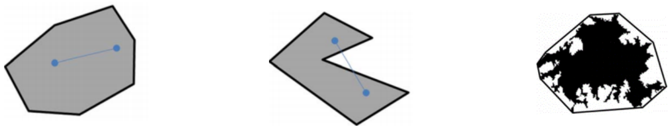

(a) (b) (c)

Supplementary Figure S. 5: Convex Hull. A cartoon of a a) convex and b) non-convex polygon. A convex polygon is a polygon which any line connecting two points inside the polygon falls inside the polygon. c. Convex hull surrounding a cell.

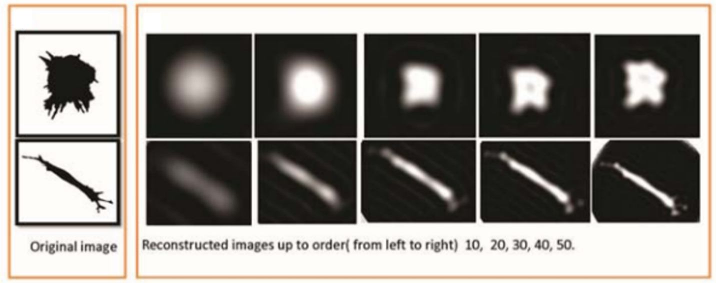


Supplementary Figure S. 6: Reconstruction of cell images for two different shaped cells. Left, original images. Right, Reconstructed images using Zernike moments up to order (from left to the right) 10, 20, 30, 40, 50.[1]

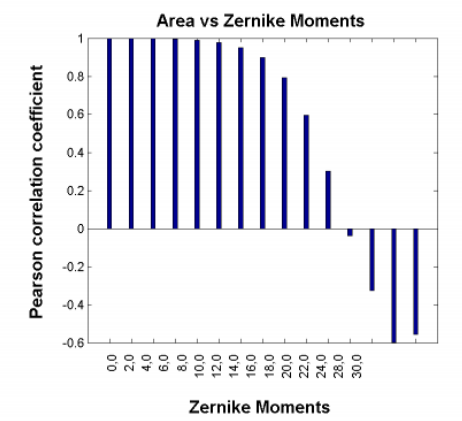


Supplementary Figure S. 7: Pearson correlation coefficient between Zernike Moments with $m = 0$ and Cell Area. Correlation is maximum and positive with Zernike moment of 0-0, it decreases with increasing the order and it is almost 0 for Zernike moment of 22-0 and it is negative for higher orders and increases in magnitude. Each data point is the average of Pearson correlation for 14 cases, 7drugs x 2 cell lines.

Supplementary Table S. 1: Description of the drugs used to perturb cytoskeleton.

| Drug | Blebbistatin | Cytochalasin D | Jasplakinolide | FAKI 14 | Nocodazole | PP2 |
|---|---|---|---|---|---|---|
| Target | Inhibits non-muscle myosin II activity | Inhibits actin polymerization | 1.Promotes actin polymerization 2.Stabilizes actin filaments | Inhibits focal adhesion kinase | Depolymerizing microtubules | Inhibits Src kinase |
| Application in cancer therapy | Inhibits cancer invasion [2] | Used as chemotherapeutic agent [3, 4]. | Reduces lung metastases of systemic Lewis lung carcinoma[5]). | Showed anti-metastatic, anti-neoplastic and anti-angiogenic properties [6, 7, 8, 9]. | Has chemotherapeutic properties by arresting cells in mitosis[10] | Prevention of cancer metastasis[11]. |
| Effect | 1.Reduces F-actin contractility 2.Increase in cell spreading and 3.Decreases cell motility[12] | Decreases cell contractility | Stabilizes F-actin [13] | Reduces cell motility and focal adhesion turnover[14]. | Cell shapes which lack tails and long protrusions and decreased cell asymmetry[15]. | Induces strong cell-cell contact and Enhances E-cadherin/catenin expression which are strongly associated with the actin cytoskeleton[11]. |
| Concentration ($\mu M$) | 10 | 1 | 0.01 | 1 | 5 | 1 |
| Vendor | Santa Cruz Biotechnology | Calbiochem | Calbiochem | Santa Cruz Biotechnology | Sigma Alrdich | Calbiochem |

Supplementary Table S. 2: Band based measurements used in the paper [16]

| Parameter | Descriptions | Formula |
|---|---|---|
| Max band intensity | Intensity of the band with maximum average intensity | Max ($I_{B_1}$, ... ,$I_{B_{10}}$ ) |
| Max band index | Location of the band with maximum average intensity | The $n$ such that $I_n$= Max ($I_{B_1}$, ... ,$I_{B_{10}}$ ) |
| Min band intensity | Intensity of the band with minimum average intensity | Min ($I_{B_1}$, ... ,$I_{B_{10}}$ ) |
| Min band index | Location of the band with minimum average intensity | The $m$ such that $I_m$= Min ($I_{B_1}$, ... ,$I_{B_{10}}$ ) |
| Above average adjusted intensity | Adjusted, weighted sum of average actin intensities greater than the mean intensity. | $\frac{\sum_{k=1}^{10} s(I_{B_k})(I_{B_k})}{Max(I_{B_1},...,I_{B_{10}})\sum_{k=1}^{10} s(I_{B_k})}$ where $s(I_{B_k}) =$ |

Supplementary Table S. 3: Gray scale measures used in this paper. See Supplementary Table S. 4 for the parameters used in this table.

| Parameter | Formula |
|---|---|
| Autocorrelation | $\sum_{i=1}^{N_g}\sum_{j=1}^{N_g} ijp(i,j)$ |
| Contrast M(Matlab) | $\sum_{i=1}^{N_g}\sum_{j=1}^{N_g} \lvert i-j\rvert^2 p(i,j)$ |
| Correlation M(Matlab) | $\sum_{i=1}^{N_g}\sum_{j=1}^{N_g} \frac{(i-\mu i)(j-\mu j)p(i,j)}{\sigma_i\sigma_j}$ |
| Correlation P | $\sum_{i=1}^{N_g}\sum_{j=1}^{N_g} \frac{ijp(i,j)-\mu_x\mu_y}{\sigma_i\sigma_j}$ |
| Cluster Standard deviation | $\sum_{i=1}^{N_g}\sum_{j=1}^{N_g} (i+j-\mu_x-\mu y)^2 p(i,j)$ |
| Cluster Prominence | $\sum_{i=1}^{N_g}\sum_{j=1}^{N_g} (i+j-\mu_x-\mu y)^4 p(i,j)$ |
| Cluster Shade | $\sum_{i=1}^{N_g}\sum_{j=1}^{N_g} (i+j-\mu_x-\mu y)^3 p(i,j)$ |
| Dissimilarity | $\sum_{i=1}^{N_g}\sum_{j=1}^{N_g} \lvert i-j\rvert p(i,j)$ |
| Energy | $\sum_{i=1}^{N_g}\sum_{j=1}^{N_g} p^2(i,j)$ |
| Entropy | $-\sum_{i=1}^{N_g}\sum_{j=1}^{N_g} p(i,j)\log(p(i,j)$ |
| Homogeneity M(Matlab) | $\sum_{i=1}^{N_g}\sum_{j=1}^{N_g} \frac{p(i,j)}{1+\lvert i-j\rvert}$ |
| Homogeneity P | $\sum_{i=1}^{N_g}\sum_{j=1}^{N_g} \frac{p(i,j)}{1+(i-j)^2}$ |
| Maximum probability | $MAX_{i,j}p(i,j)$ |
| Sum squares variance | $\sum_{i=1}^{N_g}\sum_{j=1}^{N_g} (i-\mu)^2 p(i,j)$ |
| Sum average | $\sum_{i=2}^{2N_g} ip_{x+y}(i)$ |
| Sum variance | $\sum_{i=2}^{2N_g} (i-SumEntropy)^2 p_{x+y}(i)$ |
| Sum Entropy | $-\sum_{i=2}^{2N_g} p_{x+y}(i)\log(p_{x+y}(i))$ |
| Difference Variance | $\sum_{i=0}^{N_g-1} i^2 p_{x-y}(i)$ |
| Difference Entropy | $-\sum_{i=0}^{N_g-1} p_{x-y}(i)\log(p_{x-y}(i))$ |
| Information Measure of Correlation1 | $\frac{Entropy-HXY1}{(max(HX,HY)}$ |
| Information measure Correlation 2 | $\sqrt{1-e^{-2(HXY2-HXY)}}$ |
| Inverse Difference Normalized(INN) (Matlab) | $\sum_{i=1}^{N_g}\sum_{j=1}^{N_g} \frac{p(i,j)}{1+\frac{\lvert i-j\rvert}{N_g}}$ |
| Inverse Difference Moment Normalized(Matlab) | $\sum_{i=1}^{N_g}\sum_{j=1}^{N_g} \frac{p(i,j)}{1+\frac{(i-j)^2}{N_g^2}}$ |

Supplementary Table S. 4: Parameters used in the definitions of gray scale measures in the Supplementary Table S. 3.

| Parameter | Formula |
|---|---|
| $P(i,j)$ | $ij^{th}$ entry of co-occurance probability matrix |
| $N_g$ | Highest intensity of gray level |
| $\mu$ | Mean value of $p(i,j)$ |
| $p_x(i)$ | $\sum_{j=1}^{N_g} p(i,j)$ |
| $p_y(i)$ | $\sum_{i=1}^{N_g} p(i,j)$ |
| $\mu_x$ | $\sum_{i=1}^{N_g}\sum_{j=1}^{N_g} ip(i,j)$ |
| $\mu_y$ | $\sum_{i=1}^{N_g}\sum_{j=1}^{N_g} jp(i,j)$ |
| $\sigma_x^2$ | $\sum_{i=1}^{N_g}\sum_{j=1}^{N_g} (i-\mu_x)^2 p(i,j)$ |
| $\sigma_y^2$ | $\sum_{i=1}^{N_g}\sum_{j=1}^{N_g} (j-\mu_y)^2 p(i,j)$ |
| $p_{x+y}(k)$ | $\sum_{i=1}^{N_g}\sum_{j=1}^{N_g} p(i,j)$ where $i+j=k, k=2,..,2N_g$ |
| $p_{x-y}(k)$ | $\sum_{i=1}^{N_g}\sum_{j=1}^{N_g} p(i,j)$ where $i-j=k, k=0,..,N_g-1$ |
| $HX$ | -$\sum_{i=1}^{N_g} p_x(i)\log p_x(i)$ |
| $HY$ | -$\sum_{i=1}^{N_g} p_y(i)\log p_y(i)$ |
| $HXY1$ | -$\sum_{i=1}^{N_g}\sum_{j=1}^{N_g} p(i,j)\log\left(p_x(i)p_y(j)\right)$ |
| $HXY2$ | -$\sum_{i=1}^{N_g}\sum_{j=1}^{N_g} p_x(i)p_y(j)\log\left(p_x(i)p_y(j)\right)$ |

Supplementary Table S. 5: Geometric parameters.

| Parameter | Description | Unit |
|---|---|---|
| Cell area | Area of the cell | $Pixel^2$ |
| Cell perimeter | Perimeter of the cell | $Pixel$ |
| Cell major axis | The major axis of an ellipse drawn around the cell | $Pixel$ |
| Cell minor axis | The minor axis of an ellipse drawn around the cell | $Pixel$ |
| Cell circularity | $\frac{4Area}{Perimeter^2}$ | $Unit - less$ |
| Cell aspect Ratio | $\frac{Cell\ Major\ axis}{Cell\ Minor\ axis}$ | $Unit - less$ |
| Cell roundness | $\frac{4\pi Area}{\pi\ \ Cell\ Major\ axis^2}$ | $Unit - less$ |
| Cell solidity | $\frac{Convex\ hull\ area}{Cell\ area}$ | $Unit - less$ |
| Max cell Radius | Maximum radius of the cell | $Pixel$ |
| Cell radius ratio | $\frac{Max\ radius\ of\ the\ cell}{Min\ radius\ of\ the\ cell}$ | $Unit - less$ |
| Mean Cell Radius | The mean of all radii drawn from the hull's centroid to an exterior point | $Pixel$ |
| CV Cell Radius | The relative variation of radii drawn from the cells center to an exterior point. Given by the Standard of Deviation of all Radii divided by the mean of all radii | $Unit - less$ |
| Max Span | The maximum distance from one point on the convex hull to another | $Pixel$ |

Supplementary Table S. 6: Nuclei geometric parameters.

| Parameter | Description | Unit |
|---|---|---|
| Nuc area | Area of the nucleus | $Pixel^2$ |
| Nuc perimeter | Perimeter of the nucleus | $Pixel$ |
| Nuc major | Major axis of an ellipse drawn around the nucleus | $Pixel$ |
| Nuc minor | Minor axis of an ellipse drawn around the nucleus | $Pixel$ |
| Nuc circularity | $\frac{4\pi Nuc\, area}{Nuc\, perimeter^2}$ | $Unit - less$ |
| Nuc aspect ratio | $\frac{Nuc\, major}{Nuc\, minor}$ | $Unit - less$ |
| Nuc roundness | $\frac{4\pi Nuc\, area}{\pi Nuc\, major^2}$ | $Unit - less$ |
| Nuc solidity | $\frac{Area\, of\, convex\, hull\, of\, nuclei)}{Nuc\, area}$ | $Unit - less$ |
| Distance nuc to cell centroid | Distance between the centroids of cell and nucleus | $Pixel$ |
| Nuc to cell orientation | Angle between the distance nuc to cell centroid and cell major axis | $Degree$ |

Supplementary Table S. 7: Convex hull geometric parameters.

| Parameter | Description | Unit |
|---|---|---|
| Hull area | Area of the convex hull | $Pixel^2$ |
| Hull perimeter | Perimeter of the convex hull. | $Pixel$ |
| Ratio of perimeter of Hull to cell | Ratio of hull perimeter to cell perimeter. | $Unit - less$ |
| Hull circularity | $\frac{4\pi\ Hull\ area}{Hull\ perimeter^2}$ | $Unit - less$ |
| Max hull Radius | Maximum distance from centroid of Hull to an exterior point on the hull. | $Pixel$ |
| Hull radius ratio | Maximum/minimum radius of hull. | $Unit - less$ |
| Mean hull radius | The mean of all radii drawn from the hull's centroid to an exterior point. | $Pixel$ |
| CV hull radius | (STD hull radius)/(Mean hull radius) | $Unit - less$ |
| Bounding circle diameter | The diameter of the bounding circle drawn around the cell. | $Pixel$ |
| Max circle to hull radius | The maximum distance from the center of the bounding circle to an edge of the convex hull. | $Pixel$ |
| Circle to hull radius ratio | (Max circle to hull radius)/(Min circle to hull radius) | $Unit - less$ |
| Mean circle to hull radius | The mean of all radii drawn from the circle's center to the hull | $Pixel$ |
| CV Circle2Hull Radius | (STD( all radii drawn from the circle s center to the hull ))/(Mean circle to hull radius) | $Unit - less$ |

Supplementary Table S. 8: Roughness measurements.

| Parameter | Formula | Units |
|---|---|---|
| Mean of absolute values | $R_a = \frac{1}{n}\sum_{i=1}^{n} \lvert y_i \rvert$ | *Pixel* |
| Root mean squared | $R_{rms} = \sqrt{\frac{1}{n}\sum_{i=1}^{n} y_i^2}$ | *Pixel* |
| Maximum height of the profile | $R_{yt} = R_p - R_v$ Where $R_p$ has the maximum and $R_v$ has the minimum height in the profile | *Pixel* |
| Kurtosis | $R_{Ku} = \frac{1}{nR_{rms}^4}\sum_{i=1}^{n} y_i^4$ | $Pixel^4$ |

Supplementary Table S. 9: Significance of changes in primary principal component (PPC) of morphometric classes for different drugs according to T-test analysis. The p-value of the null hypothesis that the experimental condition has the same mean as the control is reported for each class of morphometrics, based on the PPC. There are significant changes (p-value $<$ 1E-3) in some actin textural measures (Band-Based, Fractal Dimension and Gray scale measures), spreading measures (Cell Geometric, Nuclei Geometric, Hull Geometric and Zernike moment) and Irregularity measures (Waviness and Roughness) for every experimental condition. Sample size is 300 cells for each cell line and drug condition other than DLM8 FAKI 14 which has a sample size of 200.

| Comparisons: Control vs | Cell Line | Band Based | Fractal Dimension | Gray Scale | Cell Geometric | Nuclei | Hull Geometric | Zernike Moment | Waviness | Roughness |
|---|---|---|---|---|---|---|---|---|---|---|
| Blebbistatin | DUNN | 1.00E-07 | 2.60E-14 | 0.008 | 1.60E-78 | 0.006 | 3.10E-14 | 3.00E-21 | 2.00E-35 | 2.70E-45 |
| | DLM8 | 3.00E-08 | 6.30E-45 | 6.00E-09 | 6.00E-25 | 0.004 | 7.90E-08 | 0.0018 | 2.00E-12 | 5.30E-16 |
| Cytochalasin D | DUNN | 3.00E-50 | 2.10E-31 | 2.00E-23 | 3.50E-21 | 2.00E-22 | 2.50E-06 | 0.0092 | 5.00E-10 | 3.30E-27 |
| | DLM8 | 4.00E-31 | 2.10E-13 | 1.00E-23 | 4.50E-15 | 1.00E-18 | 0.05285 | 1.00E-05 | 3.00E-07 | 8.20E-07 |
| FAKI 14 | DUNN | 2.00E-09 | 5.16E-31 | 1.00E-23 | 5.20E-09 | 2.00E-04 | 0.04184 | 0.1194 | 0.1398 | 2.20E-12 |
| | DLM8 | 4.00E-04 | 6.60E-22 | 2.00E-21 | 1.50E-06 | 2.00E-06 | 0.07864 | 0.0997 | 0.3125 | 7.80E-07 |
| Jasplakinolide | DUNN | 2.00E-04 | 2.10E-13 | 9.00E-05 | 1.30E-08 | 0.001 | 0.02999 | 4.00E-05 | 0.0036 | 0.051 |
| | DLM8 | 0.003 | 2.80E-09 | 2.00E-04 | 0.00207 | 5.00E-04 | 1.20E-05 | 0.0005 | 0.0001 | 0.02117 |
| Nocodazole | DUNN | 6.00E-13 | 5.60E-31 | 3.00E-25 | 1.30E-28 | 1.00E-04 | 4.60E-10 | 9.00E-07 | 1.00E-12 | 3.30E-16 |
| | DLM8 | 0.001 | 9.80E-14 | 4.00E-07 | 1.20E-14 | 6.00E-06 | 3.00E-09 | 6.00E-14 | 3.00E-21 | 5.30E-13 |
| PP2 | DUNN | 2.00E-21 | 1.40E-11 | 2.00E-10 | 2.50E-06 | 0.003 | 0.03602 | 0.0243 | 0.0089 | 0.00387 |
| | DLM8 | 4.00E-07 | 0.00035 | 3.00E-09 | 0.00043 | 8.00E-10 | 2.70E-09 | 4.00E-13 | 4.00E-11 | 0.00078 |
